# siGMA: Modeling Label Noise and Heterogeneity in siRNA Efficacy Prediction

**DOI:** 10.64898/2026.11.20.768048

**Authors:** Aparajita Karmakar, Abdulhamid Merii, Angus Weir, Grzegorz Kudla, Mark Basham, Alex Lubbock

## Abstract

Small interfering RNAs (siRNAs) have emerged as a powerful class of therapeutics, yet identifying viable candidates remains costly and labor-intensive given the vast combination space that must be screened. This challenge is compounded by the need for chemical modifications, which are essential to mitigate off-target effects and enhance serum stability but frequently come at the expense of baseline knockdown efficacy. As a result, siRNA design requires careful, iterative optimization, further expanding the experimental burden and rendering exhaustive screening impractical. Computational prediction offers a compelling alternative, enabling rapid, low-cost identification of promising candidates prior to wet-lab validation.

In this work we present siGMA, a heteroscedastic multimodal predictive model that integrates sequence, chemical, structural, and experimental information to predict the silencing efficacy of chemically modified siRNAs. Modelling in this domain is constrained by two principal obstacles: the scarcity of experimental data and the substantial noise arising from protocol heterogeneity across publicly available datasets. We therefore propose a model that quantifies predictive uncertainty rather than issuing confident mispredictions. We calibrated the model against such noise using a combined aleatoric and epistemic uncertainty prediction. The resulting model yielded well-calibrated estimates of predictive variability on held-out folds across datasets and outperformed current state-of-the-art approaches across noisy cross-dataset conditions, including previously unseen chemical modifications and target sequences. This demonstrates our model’s ability to rank better than the baselines while rendering the noise ceiling measurable. This provides a scalable framework that could accelerate the therapeutic siRNA development pipeline while reducing the need for exhaustive empirical screening.

## Introduction

Small interfering RNAs (siRNAs) represent a powerful therapeutic modality, offering several advantages over conventional approaches. Because siRNAs act directly on messenger RNA, they can silence targets that are otherwise considered “undruggable” by small molecules or antibodies, substantially expanding the space of accessible disease targets[1].

Despite this promise, siRNAs suffer from inherent instability and a possibility of off-target effects, necessitating extensive chemical modification to achieve therapeutic viability[2]. However, the chemical space of possible modifications is vast, and screening across this space on top of screening for optimal candidate sequences is prohibitively costly and time-consuming. To address this, in silico pipelines have increasingly leveraged artificial intelligence to learn the sequence-based binding rules governing siRNA-mRNA interactions, as well as the effects of chemical modifications on potency [3],[4].

Nonetheless, several challenges continue to hinder the development of robust predictive models. Chief among these are limited data availability and the heterogeneity of experimental conditions across studies, both of which constitute major bottlenecks for model training and evaluation. Furthermore, the breadth of the chemical modification space complicates generalization to novel modifications: it is critical that models learn the functional impact conferred by the presence of a given chemistry, rather than memorizing the modification itself, as the latter would severely limit generalizability to previously unseen chemistries.

Early models in this field relied on simpler representations and architectures, such as k-mer features and one-hot encodings paired with random forest or support vector classifier models [5]. Around 2005, the field shifted toward artificial neural network approaches, coinciding with the release of the Huesken et al. dataset, at the time the largest siRNA efficacy library available [6]. Subsequently, in 2022, La Rosa et al. introduced graph-based networks to model siRNA-mRNA interactions, an approach later extended by siRNAdiscovery through enhanced feature representations and architectural refinements [7],[8]. Transformer-based models have also been applied to sequence-based prediction tasks, owing to their strength in modeling sequential dependencies, as exemplified by Oligoformer and AttSiOff [9],[10]. Notably, the majority of these models were designed for unmodified siRNAs.

Since 2024, research attention has shifted toward models capable of handling chemically modified siR-NAs. cmsiRPred introduced a multimodal architecture that represents chemical modifications via molecular fingerprints and sequences via one-hot encoding, integrating the two through a co-attention mechanism [11]. ENsiRNA subsequently extended this multimodal paradigm by incorporating a graph-based representation of siRNA-mRNA interactions [12]. More recently, a 2026 study proposed a combined multiview framework integrating graph- and transformer-based components to jointly learn from chemical modifications, sequence information, and structural features [13].

Nevertheless, current approaches remain limited in several respects. One major limitation stems from the use of leaky cross-validation schemes that inflate reported performance metrics, an issue first highlighted by siRNAdiscovery. In our prior benchmarking work, we proposed a leave-one-target-out validation scheme to enable stricter and more realistic evaluation [8], [14]. A second limitation concerns the scarcity of experimental data on modified siRNAs, which necessitates aggregating data from multiple sources, as done by the SiRNAmod and CmsiRNAdb databases [15],[16]. This aggregation, however, introduces substantial label noise arising from differences in experimental protocols and data acquisition methods across sources. Accounting for this heterogeneity is essential to building reliable models. In the present work, we therefore introduce a multimodal approach that incorporates sequence information, chemical modification data, experimental metadata and target site accessibility. Most importantly, we implement a combination of aleatoric and epistemic uncertainty to calibrate for inter-database label noise and model’s lack of knowledge due to less data. We benchmark our final model against state-of-the-art approaches by training and testing across distinct databases to assess cross-database generalization, and demonstrate that our model outperforms existing methods under this stricter evaluation regime [17].

Finally, while predictive accuracy is essential, extracting actionable design intuitions from these models is equally important for practical siRNA design. To this end, we extend our predictions on a generated dataset for generating explainability and identifying whether the design patterns from our model[18] is consistent with the actual data and known siRNA design rules.

## Materials and Methods

### Datasets

All experiments in this study were conducted using publicly available datasets. For unmodified siRNA sequences, we employed the Huesken dataset, which comprises efficacy data for siRNAs evaluated against 30 target genes[6]. For chemically modified siRNAs, we utilized the dataset generated by Bramsen et al., which includes 21 distinct chemical modification patterns evaluated against a single target, EGFP [19].

Both the Huesken and Bramsen datasets are homogeneous in origin, as each was produced under a single, controlled experimental protocol. Despite this consistency, they exhibit considerable target variability or remain sparse with respect to the chemical space explored; we used these datasets to establish baseline model performance. Real-world siRNA efficacy data, are typically generated under heterogeneous experimental conditions, reflecting variation across laboratories, protocols, and assay formats. To assess the generalizability of our model under such conditions, we additionally incorporated cmsiRNAdb, a large-scale dataset comprising over 40,000 data points spanning 36 chemical modifications and 13 target genes, aggregated from multiple independent experimental sources and research groups [16].

Given the substantial size of cmsiRNAdb, we constructed a representative 10% subset designed to preserve both the overall efficacy distribution and the relative proportion of samples contributed by each source for use in computationally intensive procedures such as ablation studies. We refer to this subset as cmsiRNA-mini.

### Pre-Processing

All datasets were preprocessed and stored in a uniform format. For each dataset, five files were generated: two modification matrices (antisense and sense), in which each row corresponds to an siRNA and each column to a nucleotide position, with “X” denoting the absence of modification and all other entries specifying the name of the modification present at that position; two corresponding nucleotide matrices (antisense and sense) following the same row–column convention; and a metadata file containing experimental annotations such as silencing efficacy, target gene, and concentration.

Due to high negative outliers in cmsiRNAdb, rows with NANs and efficacy values below 0 were excluded prior to analysis, as such outliers may bias model training and frequently reflect measurement or reporting errors.

### The Three Modalities

The performance of chemically modified siRNAs is influenced by multiple interacting factors rather than any single determinant. Accordingly, we adopted a multimodal approach in designing our model, one that jointly accounts for these multiple contributing factors rather than treating them in isolation. We have three modalities:

- The sequence level information: Sequence identity constitutes the most critical determinant of siRNA knockdown efficacy. Building on our previous work, we selected the Z-curve as the optimal sequence representation following systematic benchmarking against several alternative encoding schemes [14]. The Z-curve transforms the nucleotide sequence into a three-dimensional representation that simultaneously encodes physicochemical properties and positional context [20]. Each nucleotide is mapped to a point (*x_i_, y_i_, z_i_*) ∈ {1, 0, 1}^3^ according to three complementary base-pairing classifications (purine/pyrimidine, amino/keto, and strong/weak hydrogen bonding):

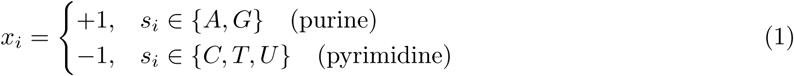

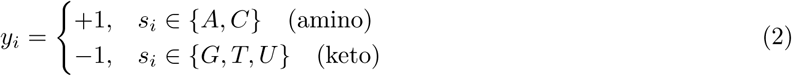

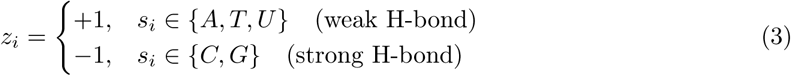

with (*x_i_, y_i_, z_i_*) = (0, 0, 0) if *s_i_* is undefined or ambiguous.

#### Locking state

Each position carries a categorical locking-state code

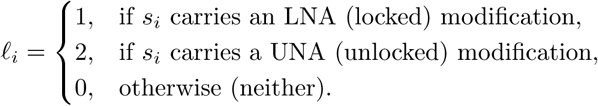

#### Modification type

Let *ϕ_i_, σ_i_, β_i_* ∈ {0, 1} be indicator variables denoting, respectively, whether nucleotide *s_i_* carries a *phosphate* modification, a *sugar* modification, or a *base* modification:

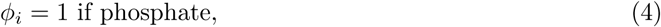

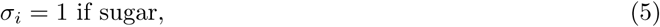

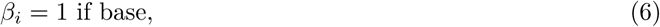

The modification-type code is the sum

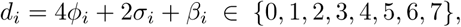

so that *d_i_* = 0 denotes *Neither*, the values {1, 2, 4} denote a single active modification class (*Base*, *Sugar*, or *Phosphate*, respectively), and any value with more than one active bit (i.e. *d_i_* ∉ {0, 1, 2, 4}) denotes a *Combined* modification. The modification type is represented as an integer, inspired by the standard integer sequence representation convention [21].

#### Full sequence representation

The complete matrix is obtained by stacking the per-position feature vectors where *N* denotes the sequence length:

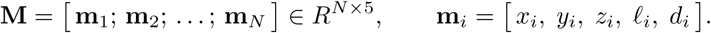

- The chemistry: Representing chemical modifications constitutes the second component of this challenge. Previous studies have approached this representation in various ways, most commonly relying on one-hot encoding or MACCS keys [22], [11]. We have implemented ErG fingerprint from [23] that encodes a molecule’s pharmacophore features (donors, acceptors, aromatics) and the distances between them, instead of atom-level structure. It finds molecules with different skeletons but similar pharmacophore patterns, which atom-based fingerprints often miss [24].

Chemical modifications were encoded as a matrix **M** ∈ *R^N^*^×^*^F^*, where *N* denotes the sequence length and *F* denotes the fingerprint length, such that each row **m***_i_* ∈ *R^F^* (*i* = 1*,…, N*) represents the fingerprint vector encoding the chemical modification at position *i* of the sequence.

- This modality encodes two categories of information: (1) experimental parameters, including concentration, oligonucleotide length, and transfection time; and (2) target site accessibility. Experimental factors are frequently excluded from activity prediction models, despite often being primary drivers of observed knockdown. Target site accessibility, determined by local mRNA secondary structure, is equally critical, as it dictates whether effective binding can occur at all. To quantify this, per-nucleotide base-pairing probabilities were computed with RNAplfold (python ViennaRNA 2.7.2) and the mRNA binding region was used with 20 flanking bases on either side of the binding region. These probabilities were averaged separately over the full binding region and the seed region, as seed region is more influential given its greater contribution to silencing efficacy [25]. The sequence-level metadata is represented as a 1 × 5 feature vector

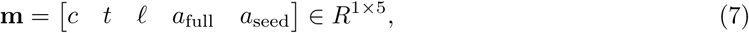

where *c* denotes the *concentration*, *t* denotes the *transfection time*, *ℓ* denotes the sequence *length*, *a*_full_ denotes the *mRNA full binding accessibility*, and *a*_seed_ denotes the *mRNA seed accessibility*.

### Multi-Modal Model

#### Model components

- Row-encoder: Every modality goes through a row encoder that goes over each nucleotide’s sequence/chemistry representation to convert it into a common embedding size. For the sequence and chemistry modalities 1 dimensional convolution neural network is used and 1D MLP is used for metadata.

Modalities *m* ∈ {seq, chem, meta} provides a per-nucleotide input 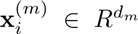 for positions *i* = 1*,…, N*. Meta modality is a single vector input. A modality-specific row-encoder 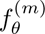 maps each position into a shared embedding space:

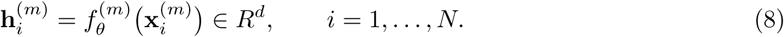

For the sequence and chemistry modalities, 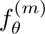 is a 1D convolutional network applied along the nucleotide axis, capturing local context around each position. For the metadata modality, 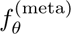 is a 1D MLP applied independently to each row.

- Row-relationship Encoder: Following the embedding stage, we introduce a Row-Relationship Encoder to model dependencies among nucleotides across the entire siRNA sequence. Whereas the embedding layer produces a local context level representation for each nucleotide, biological function often depends on how nucleotides interact across the full length of the strand interactions that are not necessarily local or sequential.

To capture these relationships, we employ a Transformer encoder block built on the self-attention mechanism. Unlike recurrent architectures such as LSTMs or bidirectional LSTMs, which process nucleotides sequentially and propagate information step-by-step, self-attention allows every nucleotide to directly attend to every other nucleotide in the sequence in a single operation. This enables the model to capture both local motifs and long-range positional dependencies simultaneously.

- Cross-attention: Once all intra-sequence relationships have been learned by the self-attention encoder, a cross-attention module is introduced to allow the chemical representation to interact with the sequence representation, thereby capturing the mutual dependencies between the two modalities.

#### Formulation

Let **H***_s_* ∈ *R^N^*^×*d*^ denote the sequence embeddings and **H***_c_* ∈ *R^N^*^×*d*^ denote the chemistry embeddings, where *N* is the sequence length and *d* is the shared hidden dimension.

The cross-attention query is derived from the chemistry embeddings, while the keys and values are derived from the sequence embeddings:

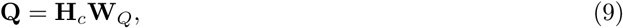

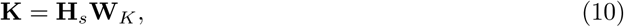

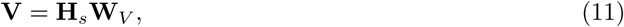

where **W***_Q_,* **W***_K_,* **W***_V_* ∈ *R^d^*^×*dk*^ are learnable projection matrices.

The cross-modal attention output is then computed as

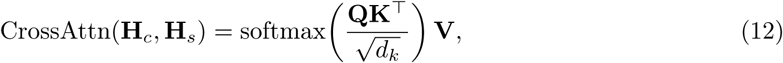

which produces an updated chemistry representation 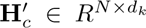, in which each chemical token attends over all sequence tokens, capturing the dependency of chemistry on sequence context (and, symmetrically, the reverse direction is computed by swapping the roles of **H***_c_* and **H***_s_*). Our model uses cross attention on both directions (chemistry to sequence and vice versa)

- Attention-pooling: Following cross-attention, an attention pooling layer is used to aggregate the token-level representations into a single fixed-length vector. Rather than applying simple mean or max pooling, the model learns an importance weight *α_i_* for each token, allowing it to emphasize the most informative positions while suppressing less relevant ones. The pooled representation is obtained as a weighted sum of the token embeddings,

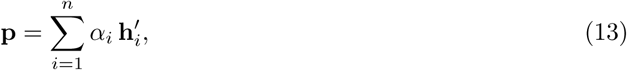

where *α_i_* denotes the learned weights. This yields a compact summary vector that retains the most salient cross-modal information for downstream prediction.

- Head: The pooled embeddings from sequence and chemistry, denoted **e**_1_ ∈ *R^d^* and **e**_2_ ∈ *R^d^* respectively, are stacked together with the metadata embedding **e***_m_* ∈ *R^d^* to form a joint representation

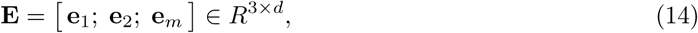

which is aggregated via mean pooling across the modality axis,

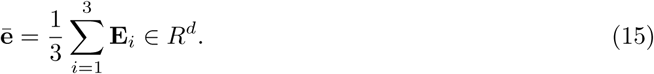

The pooled vector **ē** is passed to a regression head consisting of a shared hidden layer followed by two parallel linear branches, which emit the predicted mean *ŷ* and the predicted standard deviation *σ̂*. The latter is passed through a softplus transform to constrain it to positive values, yielding a per-sample uncertainty estimate alongside each point prediction.

### Heteroscedastic Noise

#### Aleatoric Uncertainty via Heteroscedastic Loss

Experimental siRNA knockdown measurements exhibit substantial heterogeneity arising from differences in cell lines, transfection protocols, assay platforms, and measurement noise. To explicitly account for this inherent variability, we model aleatoric uncertainty directly in the training objective [26]. Rather than producing a single point estimate, the network is trained to output both a predictive mean efficacy *µ̂* (*x_i_*) and a predictive standard deviation *σ̂* (*x_i_*) for each input *x_i_*, thereby capturing input-dependent (heteroscedastic) noise.

Under a Gaussian observation model, *y_i ∼_ N(µ̂* (*x_i_*)*, σ̂*^2^(*x_i_*)), the network parameters are optimised by minimising the negative log-likelihood (NLL) of the observed targets. The aleatoric loss is given by

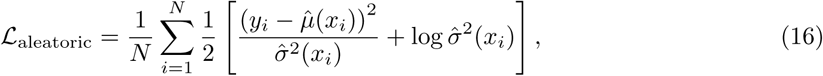

where *N* denotes the number of training samples, *y_i_*is the observed efficacy, and *µ̂* (*x_i_*) and *σ̂*^2^(*x_i_*) are the predicted mean and variance, respectively.

The two terms in Eq. 16 act in opposition. The first term penalises prediction errors in proportion to the inverse of the predicted variance. The model can therefore attenuate the influence of samples it deems noisy by assigning them a larger *σ̂*^2^(*x_i_*). The second term penalises large predicted variances and prevents the trivial solution of declaring every sample highly uncertain. The resulting balance encourages the network to assign low variance where its predictions are accurate and high variance where the observations are intrinsically variable. As a result, *σ̂*(*x_i_*) serves as a calibrated, input-specific estimate of the irreducible measurement noise associated with each siRNA.

#### Epistemic Uncertainty via Deep Ensembles

While the heteroscedastic loss in Eq. 16 captures noise intrinsic to the observations, it does not reflect the model’s uncertainty arising from limited or unevenly distributed training data. Such epistemic uncertainty is particularly relevant for siRNA efficacy prediction, where publicly available datasets are relatively small and sparsely cover the space of possible sequences and chemical modifications [27]. To estimate this component, we employ a deep ensemble of *M* = 5 independently trained models. Each member shares the same architecture and training objective but is initialised with a different random seed.

For a given input *x*, each ensemble member *m* ∈ {1*,…, M*} produces a predictive mean *µ̂_m_*(*x*) and variance 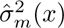. The ensemble prediction is obtained by treating the members as a uniformly weighted Gaussian mixture, whose mean is

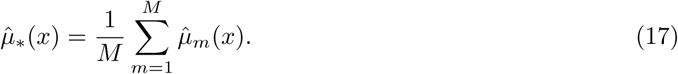

The total predictive variance then decomposes into aleatoric and epistemic components according to the law of total variance:

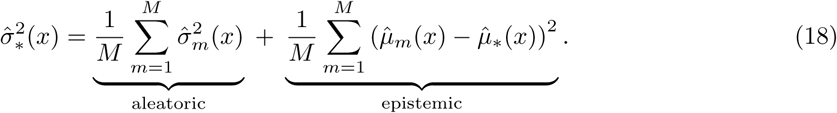

The first term, the average of the members’ predicted variances, quantifies the expected observation noise at *x*. The second term, the variance of the members’ mean predictions, captures their disagreement. Disagreement is expected to be small in densely sampled regions of the input space and large for inputs that lie far from the training distribution. We accordingly report the epistemic standard deviation *σ̂*_epi_(*x*), defined as the square root of the second term in Eq. 18, as an indicator of insufficient knowledge attributable to sparse or underrepresented training data. The aleatoric standard deviation *σ̂*_ale_(*x*), defined analogously from the first term, reflects irreducible measurement variability. Reporting the two components separately allows distinguishing between predictions that are uncertain because the underlying measurements are inherently noisy from those that are uncertain because the model has seen few comparable examples. Only the latter can be reduced by acquiring additional training data.

### Choice of Cross-Validation

As established in our previous work, the choice of cross-validation strategy is a critical methodological consideration, since conventional approaches such as holdout and k-fold cross-validation can introduce data leakage when structurally or chemically similar sequences are distributed across both training and test partitions, thereby inflating apparent model performance. To mitigate this risk, we employed dataset-specific, leakage-aware cross-validation schemes tailored to the structure of each dataset used in this study.

For the Huesken dataset, we implemented leave-one-target-out cross-validation, whereby all siRNA sequences directed against a given mRNA target were withheld as a single fold, ensuring that no target-specific sequence information was shared between training and test sets. For the Bramsen dataset, which comprises siRNAs bearing diverse chemical modification patterns, we adopted a leave-one-modification-out scheme, excluding all sequences carrying a particular modification type from the training set in each fold, thereby preventing modification-specific bias from leaking across partitions.

Besides the above two, for the cmsiRNAdb dataset, we additionally tested a leave-one-source-group-out cross-validation strategy, in which data sources were withheld as complete folds. Given that the number of sequences contributed by individual sources was often too small to constitute a statistically meaningful and balanced fold, we aggregated sources at the level of their region/patent type (WO – World Intellectual Property Organization, US – United States, CN – China and TW – Taiwan), forming larger, more balanced source groups. This aggregation reduced class imbalance that would otherwise arise from highly heterogeneous per-source sample sizes, while preserving the underlying rationale of source-level generalization assessment as the same sources would not occur in the train and test. cmsiRNAdb is a database curated from multiple sources. We evaluated model performance across different group splits to determine whether results remain consistent. Although some sequence regions may overlap between datasets from different laboratories, the purpose of this experiment was solely to assess whether the model can generalize to unseen targets and chemistries.

Collectively, these dataset-specific cross-validation schemes were designed to provide a more rigorous and realistic estimate of model generalizability by explicitly preventing leakage of related sequences, chemistries, or setup information between training and evaluation partitions.

### Comparison with State of the Art Models

To rigorously assess generalization performance, we benchmarked our model against two state-of-the-art baselines, ENsiRNA [12] and cmsiRPred [11]. All models were trained on cmsiRNAdb and subsequently evaluated on the independent Bramsen dataset, which was deliberately chosen because it introduces chemical modifications, target sequences, and experimental conditions that are absent from the training distribution, thereby constituting a stringent test of out-of-distribution generalization. We note, however, that this comparison is not entirely like-for-like: our model is explicitly formulated to calibrate for experimental noise, whereas the baseline methods are not, and any difference in performance should therefore be interpreted with this distinction in mind.

### Evaluation Metrics

Let 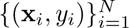 denote the test set, where *y_i_* is the observed value and *ŷ_i_* is the model prediction. For each prediction the model also provides an aleatoric standard deviation *σ_a,i_* and an epistemic standard deviation *σ_e,i_*. Assuming the two sources are independent, their variances add, and the total predictive standard deviation is

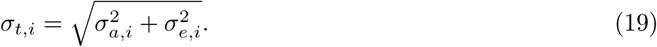

We evaluate both predictive accuracy and uncertainty quality using the following metrics.

**1. Mean Squared Error (MSE):**

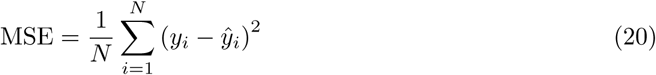

**2. Root Mean Squared Error (RMSE):**

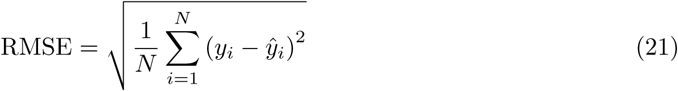

**3. Area Under the ROC Curve (ROC-AUC):** The continuous targets are binarised with a threshold *τ*, so *c_i_* = 1[*y_i_ τ*], and *ŷ_i_* is used as the ranking score. With the decision threshold *t*, the true and false positive rates are

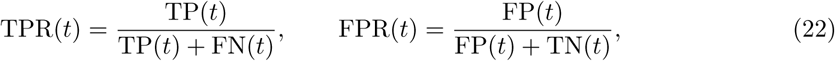

and

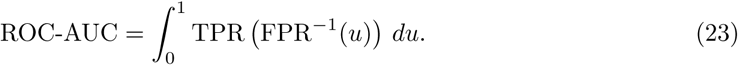

**4. Area Under the Precision–Recall Curve (PR-AUC):**

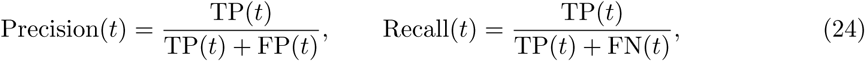

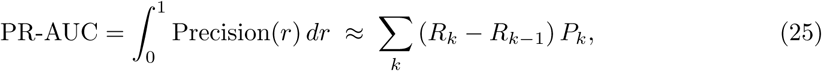

where *P_k_*and *R_k_* are precision and recall at the *k*-th threshold.

5. **Spearman Rank Correlation (***ρ*): With *d_i_* = rank(*y_i_*) − rank(*ŷ_i_*) (no ties),

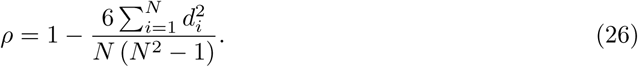

In the presence of ties, *ρ* is computed as the Pearson correlation between the rank variables.

**6. Average Predictive Standard Deviation (Avg SD):**

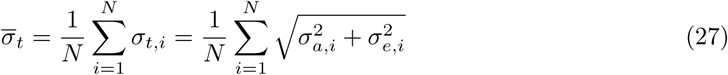

7. **SD-to-RMSE Ratio:** This checks whether the predicted uncertainty matches the observed error magnitude:

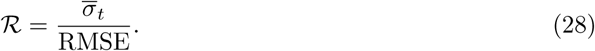

*R* ≈ 1 indicates well-calibrated uncertainty; *R <* 1 indicates over-confidence and *R >* 1 under-confidence.

**8. 90% Confidence Interval Coverage:** Assuming a Gaussian predictive distribution, the 90% interval is

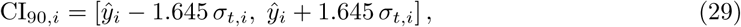

and the empirical coverage is

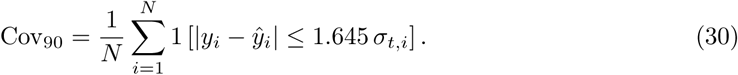

A well-calibrated model gives Cov_90_ ≈ 0.90.

**9. Aleatoric Fraction of Total Uncertainty:**

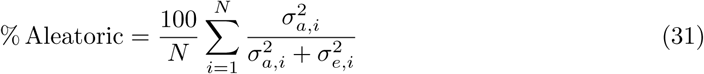

**10. Epistemic Fraction of Total Uncertainty:**

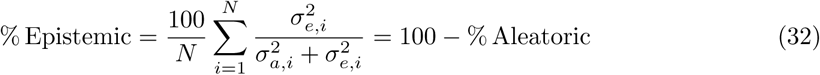

### Identifying Design Patterns

Beyond establishing overall predictive performance and robustness, we tested whether our model captures and rediscovers the underlying modification design patterns previously established through wet-lab experiments. Using the set of modifications *C* reported in the dataset of [19], we generated a dummy dataset by systematically placing each modification at every position of the sequence, one at a time:

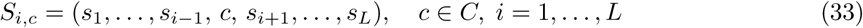

where *S* is the reference sequence of length *L*. We used the unmodified sequence common to the entire Bramsen dataset, since the model is trained on various modifications of this one sequence. This produced an exhaustive panel covering every position–modification combination. We then used our model, trained on the Bramsen dataset, to predict outcomes for each variant in this dummy set and compared the resulting positional preferences against the modification patterns reported in the original paper, to assess how well the model’s learned behavior aligns with established experimental findings.

### Experimental Setup

All models were trained on a single NVIDIA Quadro RTX 4000 GPU (8 GB VRAM) for 100 epochs. While the outer cross-validation used leave-one-group-out, we also applied group-based splitting for internal validation; a single GroupShuffleSplit ensured that no group appeared in both the training and validation sets. The checkpoint with the best performance on this internal validation split was kept for evaluation. In the cross-validation experiments, the learning rate was set to 10^−4^. For the comparison with state-of-the-art methods, all three models were trained under the same protocol (100 epochs, best-checkpoint selection) with a learning rate of 10^−4^. ENsiRNA and cmsiRPred has been trained using the default hyperparameters. Training and evaluation were performed with a batch size of 128, and 15% of the training data was held out fo the validation set. A fixed classification threshold of 0.5 was used throughout for ROC-AUC computation, since several datasets and subgroups exhibit strongly imbalanced class distributions, for which higher thresholds would yield undefined (NaN) values. Given the limited size of the dataset a compact model dimension of 64 was used. The encoded sequences were then processed by a Transformer encoder with 2 layers, each with 2 attention heads and a feed-forward dimension of 128, and a dropout rate of 0.2 was applied throughout.

## Results and Discussion

### Model Performance across Datasets

Table 1 compares model performance across all datasets. For each dataset we report empirical coverage at the 90% confidence interval, the ratio of RMSE to the mean predicted standard deviation, ROC-AUC, PR-AUC and the decomposition of predictive variance into aleatoric and epistemic components.

**Table 1:**
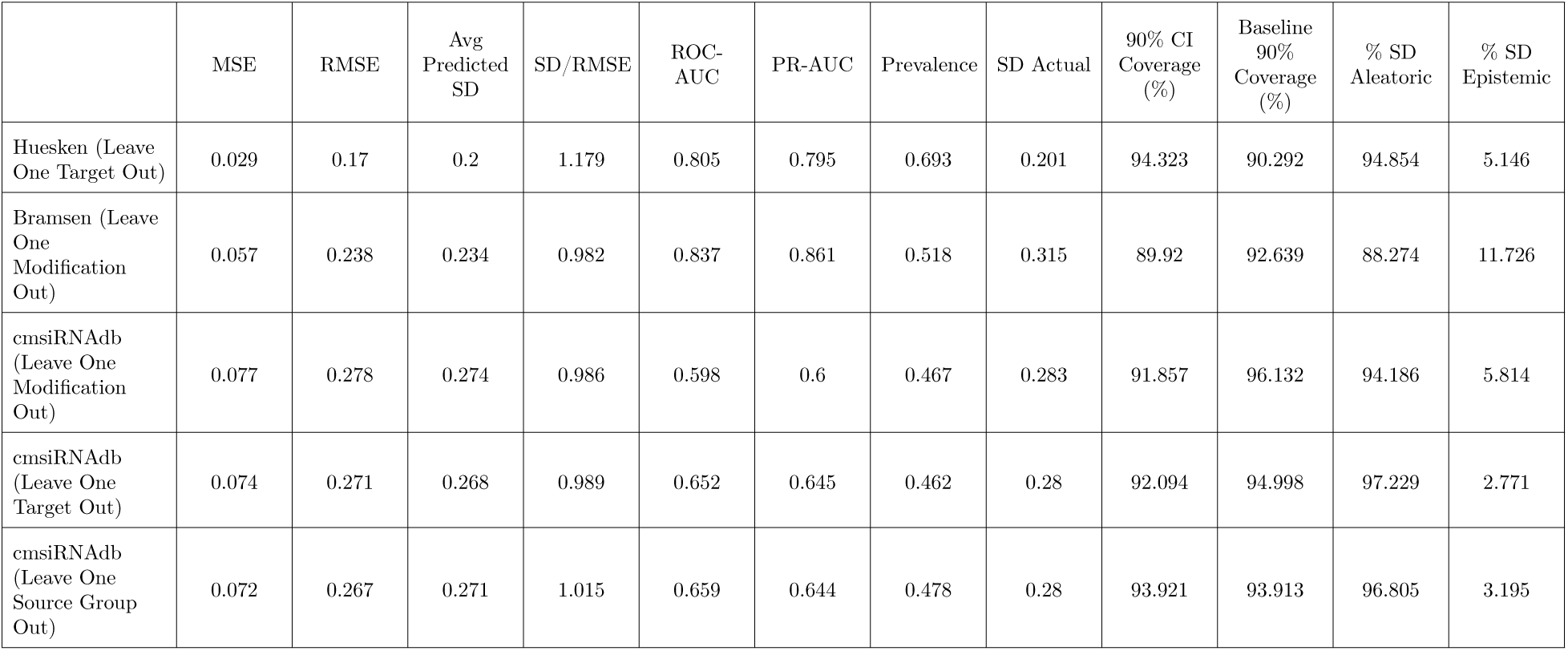
Cross-Validation metric summary across datasets showing Prediction errors, Prediction Coverage and Uncertainty decomposition.

|  | MSE | RMSE | Avg Predicted SD | SD/RMSE | ROC-AUC | PR-AUC | Prevalence | SD Actual | 90% CI Coverage (%) | Baseline 90% Coverage (%) | % SD Aleatoric | % SD Epistemic |
|---|---|---|---|---|---|---|---|---|---|---|---|---|
| Huesken (Leave One Target Out) | 0.029 | 0.17 | 0.2 | 1.179 | 0.805 | 0.795 | 0.693 | 0.201 | 94.323 | 90.292 | 94.854 | 5.146 |
| Bramsen (Leave One Modification Out) | 0.057 | 0.238 | 0.234 | 0.982 | 0.837 | 0.861 | 0.518 | 0.315 | 89.92 | 92.639 | 88.274 | 11.726 |
| cmsiRNAdb (Leave One Modification Out) | 0.077 | 0.278 | 0.274 | 0.986 | 0.598 | 0.6 | 0.467 | 0.283 | 91.857 | 96.132 | 94.186 | 5.814 |
| cmsiRNAdb (Leave One Target Out) | 0.074 | 0.271 | 0.268 | 0.989 | 0.652 | 0.645 | 0.462 | 0.28 | 92.094 | 94.998 | 97.229 | 2.771 |
| cmsiRNAdb (Leave One Source Group Out) | 0.072 | 0.267 | 0.271 | 1.015 | 0.659 | 0.644 | 0.478 | 0.28 | 93.921 | 93.913 | 96.805 | 3.195 |

Coverage is close to nominal in every dataset, with Huesken et al. showing slightly optimistic behaviour (empirical coverage above the nominal level at 90%). The RMSE/SD ratio is near unity throughout, indicating that the magnitude of the predicted uncertainty tracks the magnitude of the actual error rather than being systematically inflated or deflated. The uncertainty estimates are therefore well calibrated in both the interval and the scale sense.

Mean squared error increases systematically with dataset difficulty. The Huesken dataset, comprising unmodified siRNAs from a single experimental source, yields the lowest MSE and the narrowest standard deviations, whereas the poorest performance is observed on cmsiRNAdb, which aggregates heterogeneous sources and carries substantially higher label noise.

The calibration result is most informative where point accuracy is weakest. On cmsiRNAdb the model achieves high MSE and low ROC-AUC, yet the predicted standard deviations remain well calibrated: the model is accurate about how inaccurate it is. To assess how well our model captures predictive uncertainty, we compared it against a naive baseline that predicts only the training-set mean and standard deviation for every sample. Our model’s empirical coverage was closer to the nominal level in most settings while the baseline was much higher. The exceptions were the Huesken dataset and the leave-one-source-group-out split for cmsiRNAdb, where our model over-covered (produced wider-than-nominal intervals), one possibility owing to the fewer Source Groups in cmsiRNAdb. This separation between accuracy and calibration indicates that the loss on this dataset is not a failure of the learned mapping but a property of the labels themselves, and that explicit noise modelling is an essential rather than optional component when working with sparsely characterised, experimentally heterogeneous data.

The variance decomposition in Figure 3 supports this interpretation. Aleatoric variance consistently dominates epistemic variance across datasets, meaning that most of the predictive uncertainty is irreducible by additional modelling and is attributable to label noise. The exception is Bramsen et al., where the epistemic contribution is comparatively larger, consistent with sparse experimental sampling of a vast chemical space rather than with noisy measurement of the regions that are sampled.

**Figure 1:**
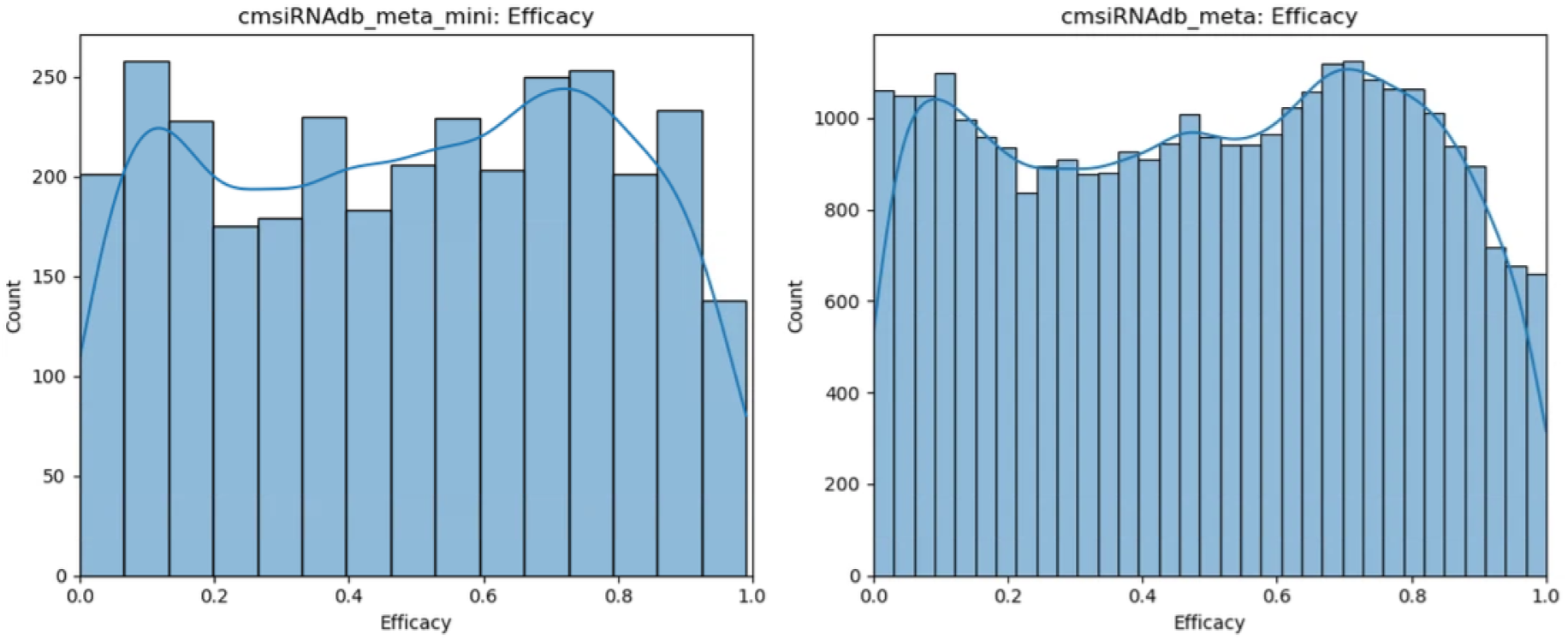
Efficacy distribution across cmsiRNAdb and cmsiRNA-mini.

**Figure 2:**
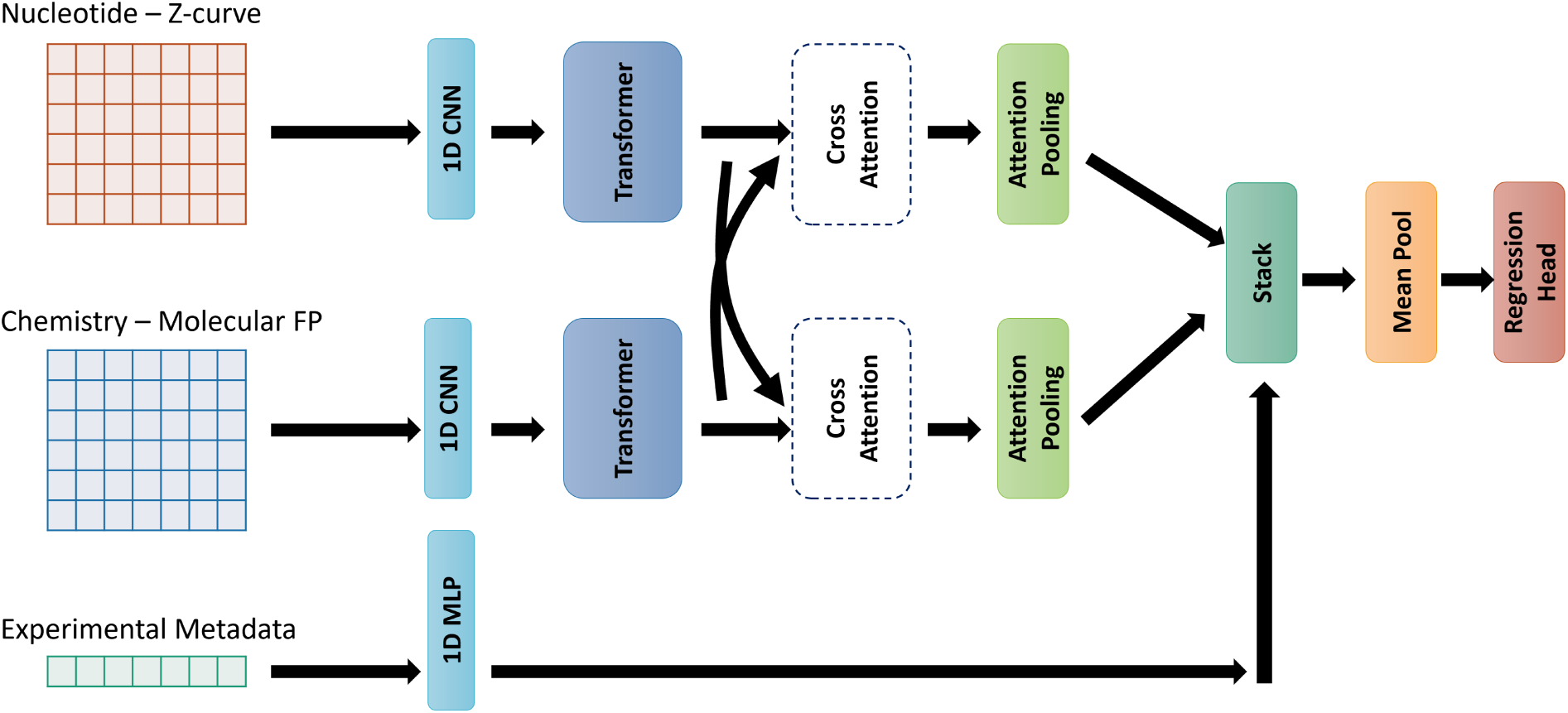
Multimodal architecture of our model.

**Figure 3:**
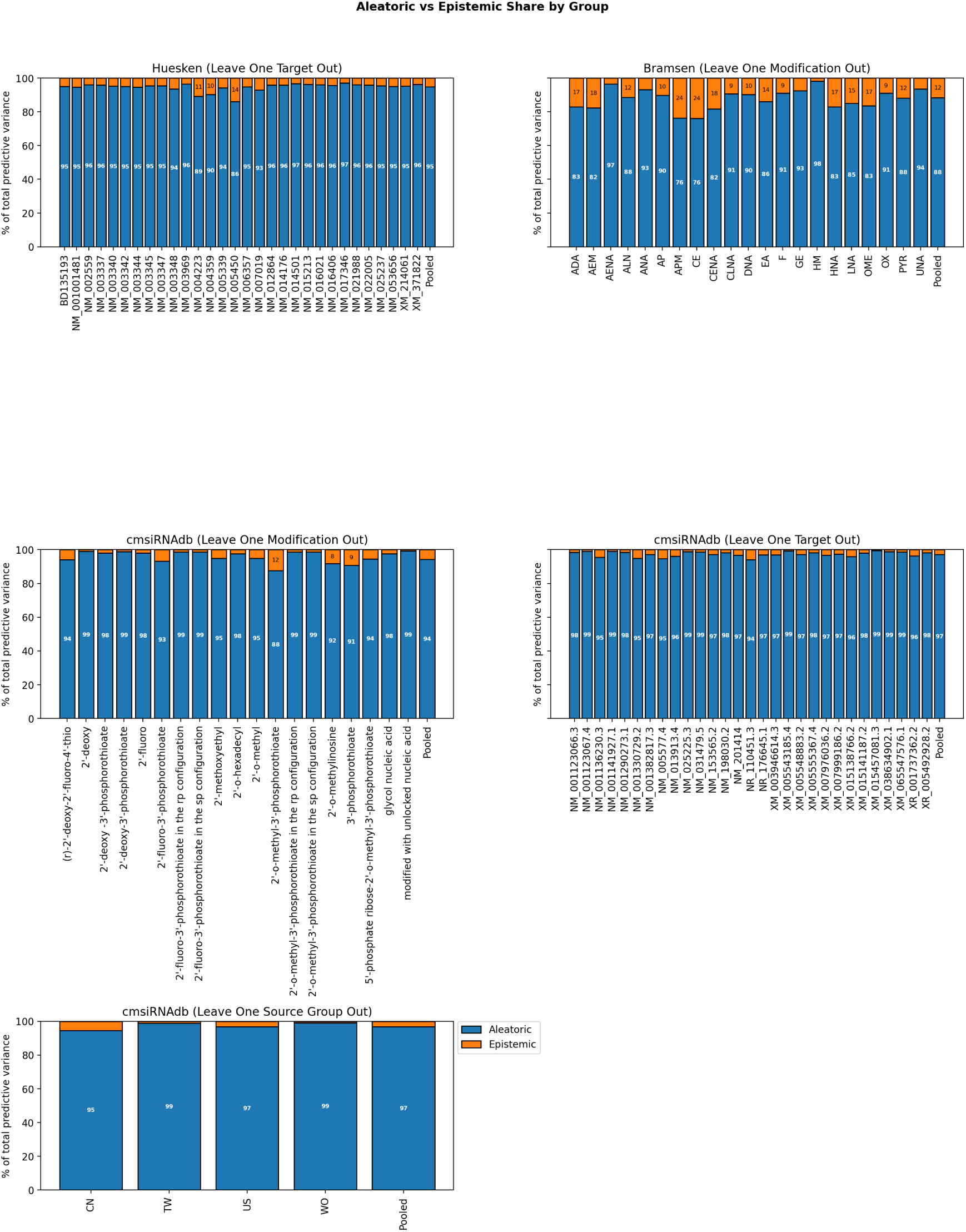
The decomposition of aleatoric and epistemic contributions show high aleatoric noise which points towards high label noise in the data rather than lack of knowledge due to limited examples.

A second structural issue is visible in the grouped analysis. Each metric (MSE, Spearman, ROC-AUC and PR-AUC) in Figure 4, 5 and 6 computed group by group vary widely. Aggregate metrics computed over a pooled test set therefore conflate two different quantities and tend to report a performance level that is not attainable on an unseen group. Standard k-fold cross-validation does not expose this, since random folds place members of the same group on both sides of the split and the reported score reflects interpolation within groups rather than generalisation across them. Grouped or scaffold-aware splitting is required for the reported numbers to carry their intended meaning.

**Figure 4:**
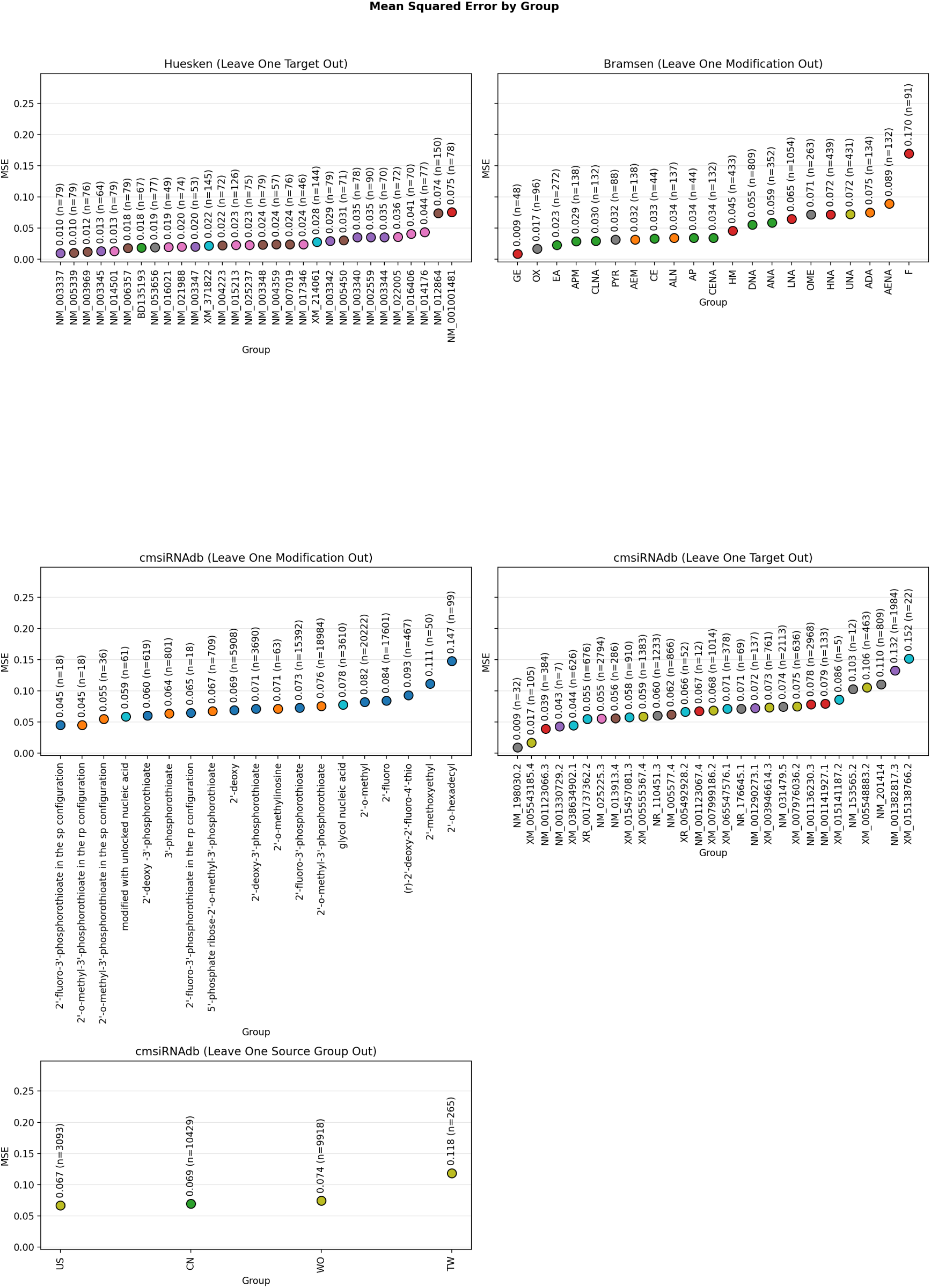
Mean square errors of the average point estimation of efficacy across groups in each dataset.

**Figure 5:**
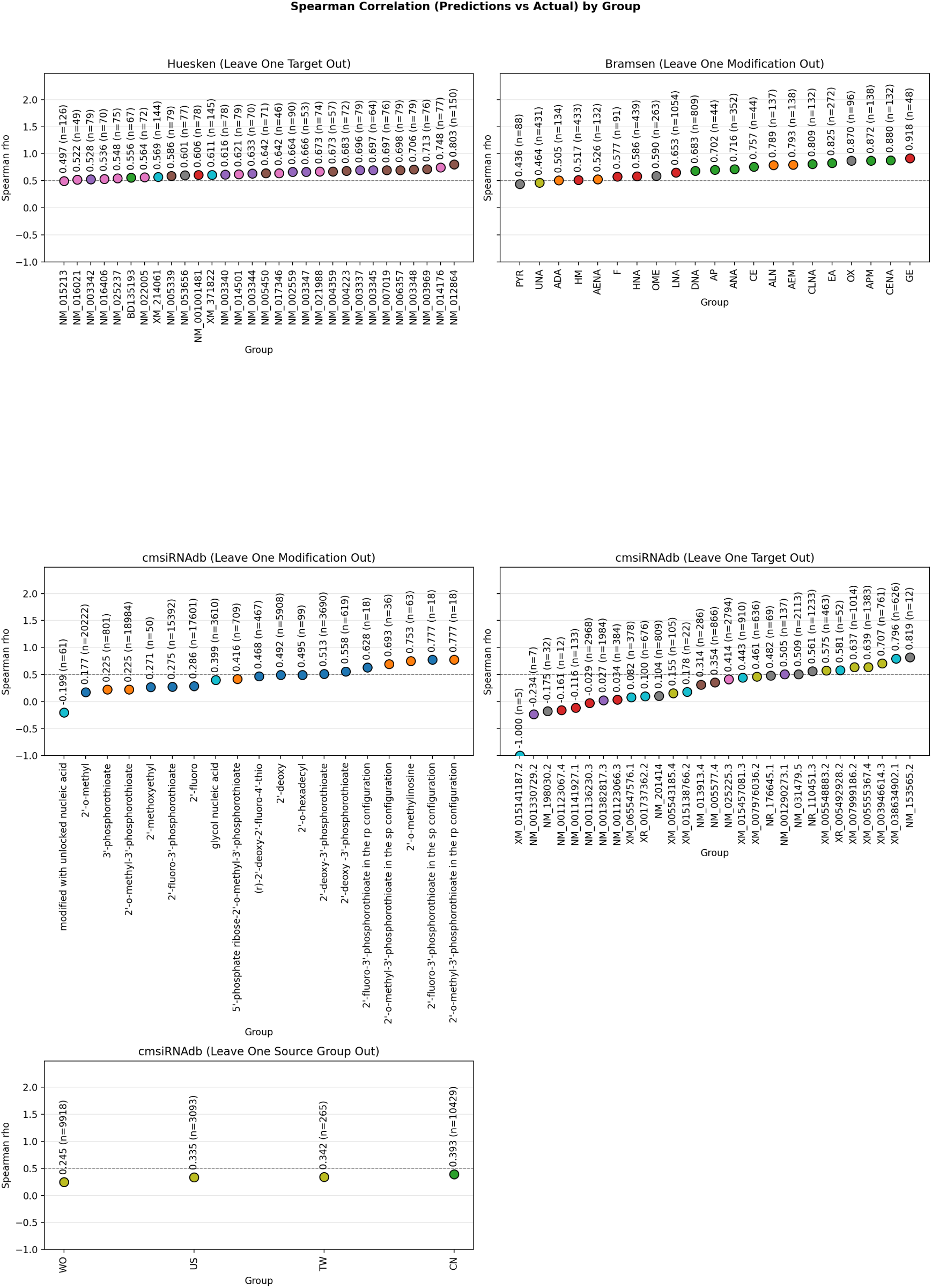
Spearman correlation across groups for different datasets show a lot of variations from group to group with lower performances across cmsiRNAdb.

**Figure 6:**
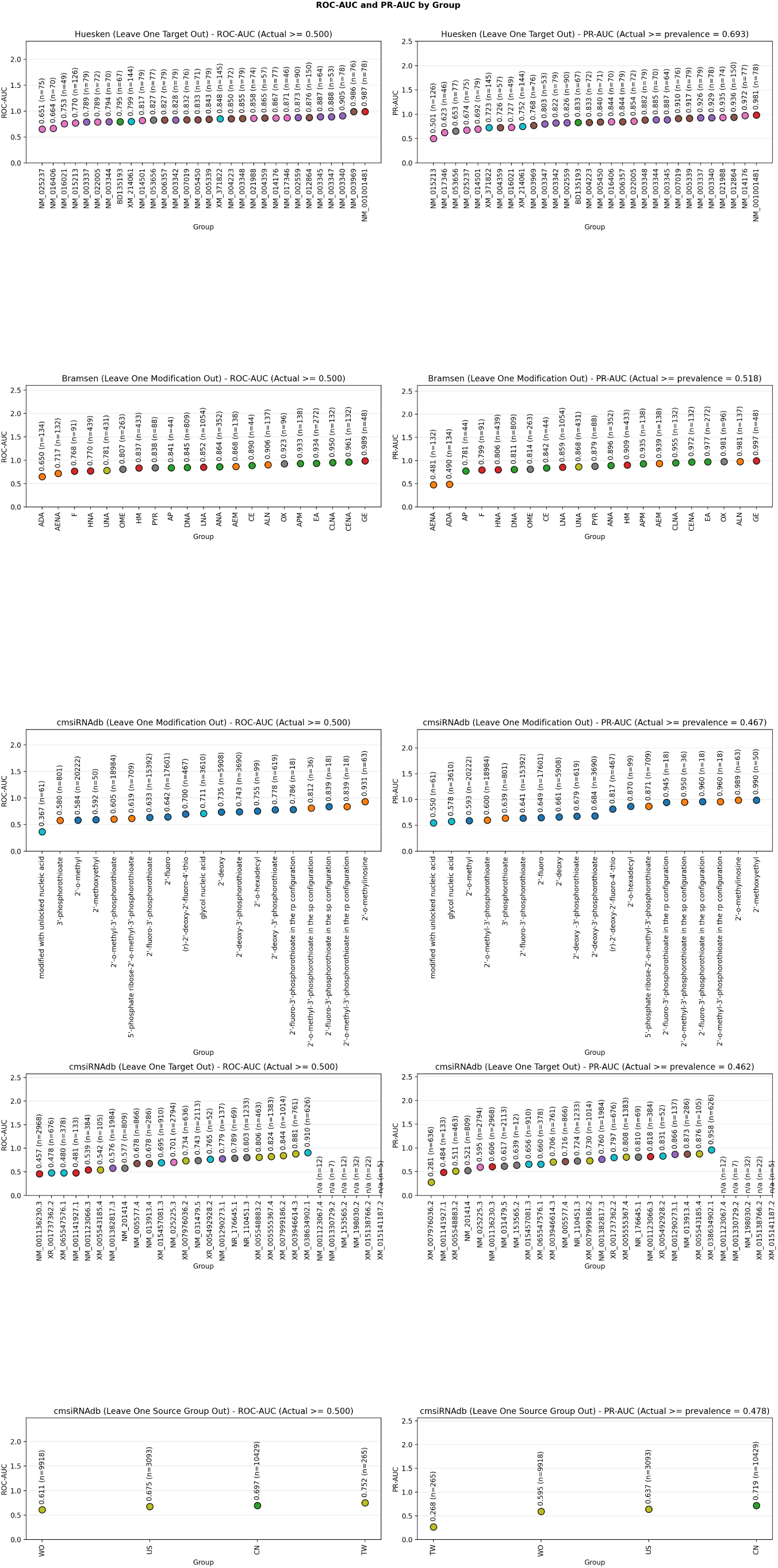
PR-AUC and ROC-AUC across groups fo1r5 each dataset at a 0.5 threshold for ROC-AUC and prevalance for PR-AUC shows consistently higher performances across groups in Huesken and Bramsen datasets which drops for cmsiRNAdb.

Taken together, the high and consistent aleatoric fraction, the group-dependent spread in metrics, and the combination of poor accuracy with well-calibrated uncertainty on cmsiRNAdb raise a question that precedes model selection: whether these datasets carry enough reliable signal to support the conclusions commonly drawn from them. Well-calibrated uncertainty makes the limitation measurable rather than removing it; the calibrated standard deviations should be read as an estimate of the noise ceiling these datasets impose on any model trained on them.

### Comparison with State of the Art Models

As noted above, a direct comparison is complicated by the fact that existing models are calibrated toward different objectives. Nevertheless, we benchmarked our model against two state-of-the-art methods, cm-siRPred and ENsiRNA. All models were trained on the cmsiRNAdb dataset, which carries substantial label noise, and evaluated on the Bramsen dataset. Under this setting (Figure 7), our model achieved a 17.7% lower MSE, 185.6% increase in Spearman correlation, 25.5% improvement in ROC-AUC and 29.9% improvement in PR-AUC than ENsiRNA. It also gained 1.4% improvement on MSE, 291.2% gain on Spearman, 30.2% and 27% improvement in AUCs over cmsiRPred. This pattern indicates that although absolute point-estimate accuracy is limited by the inherent noise in the training labels leading to high MSE and low Spearman, the model retains considerably stronger ranking ability. By jointly modelling sequence, chemistry and experimental metadata and by disentangling aleatoric from epistemic uncertainty the former through explicit calibration, the latter through ensembling, siGMA surpasses current state-of-the-art methods, which model the efficacy outcome in isolation and leave the underlying noise unaccounted for.

**Figure 7:**
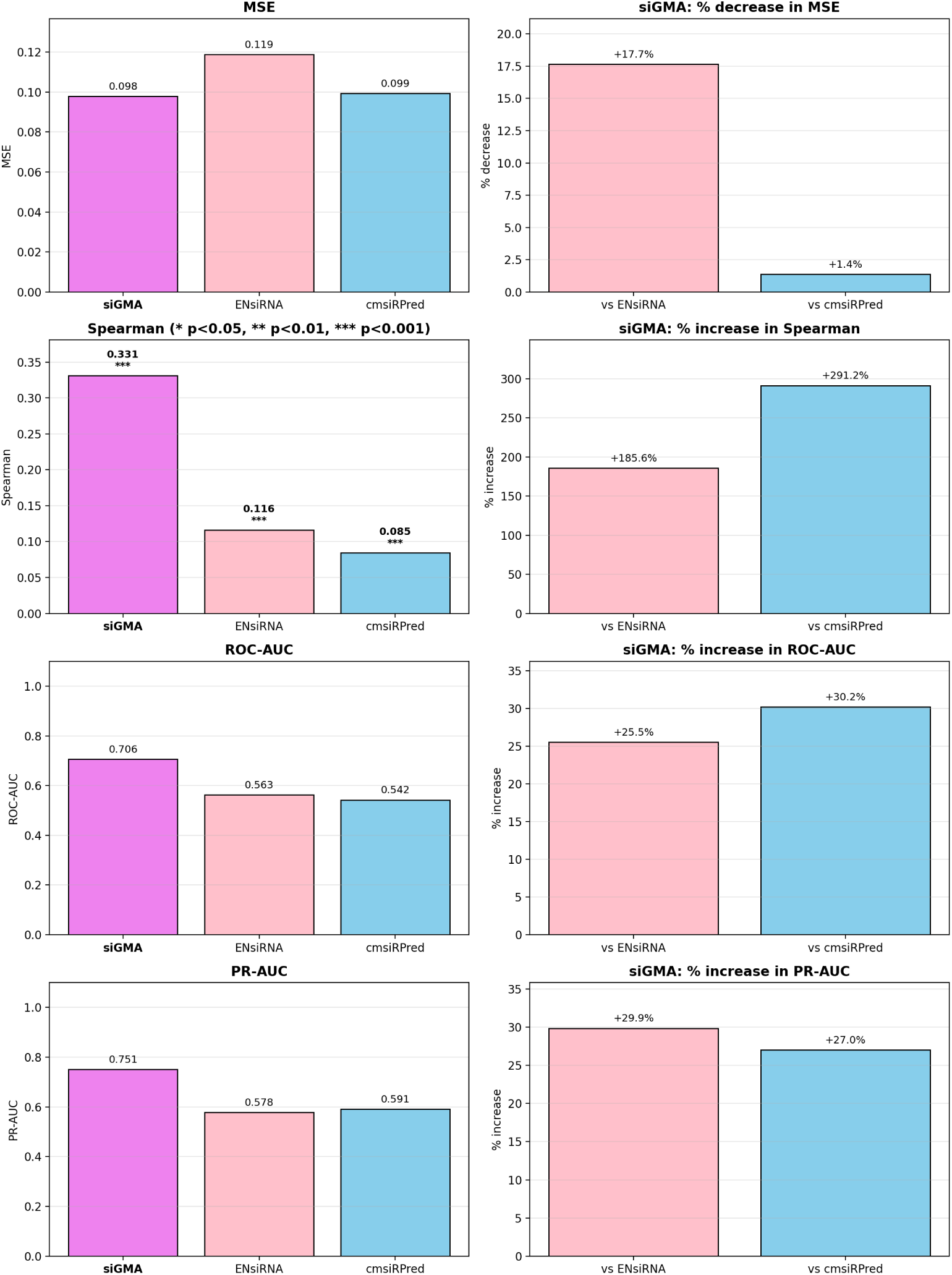
Comparison with the State of the Art model shows our model siGMA outperforms the rest in all metrics. We have compared MSE, Spearman and it’s significance, ROC-AUC and PR-AUC across the three models as well as demonstrated the percentage increase/decrease in metrics attained by siGMA.

### Design Patterns Uncovered by the Model

To evaluate our model’s ability to find meaningful design patterns that is in the data, we trained it on Bramsen’s experimental dataset and generated predictions on a synthetic dataset constructed by systematically placing each modification at each position along both strands. This analysis revealed (Figure 8) that 3^′^ overhangs of the antisense strand improve or retain potency of the unmodified baseline (X_X-0.78), the sense strand tolerates nearly all modifications well, and the 5^′^ end and seed region of the antisense strand are highly sensitive to modifications, particularly bulky ones. These findings are in strong agreement with Bramsen’s wet-lab observations, providing validation of the model’s ability to learn binding-relevant patterns from a dataset which is already sparse. These are, of course, patterns present in the training data, but a model’s ability to recover them is a prerequisite for trusting it: a model that fails to reproduce well-established design rules offers little confidence when extrapolating to untested designs.

**Figure 8:**
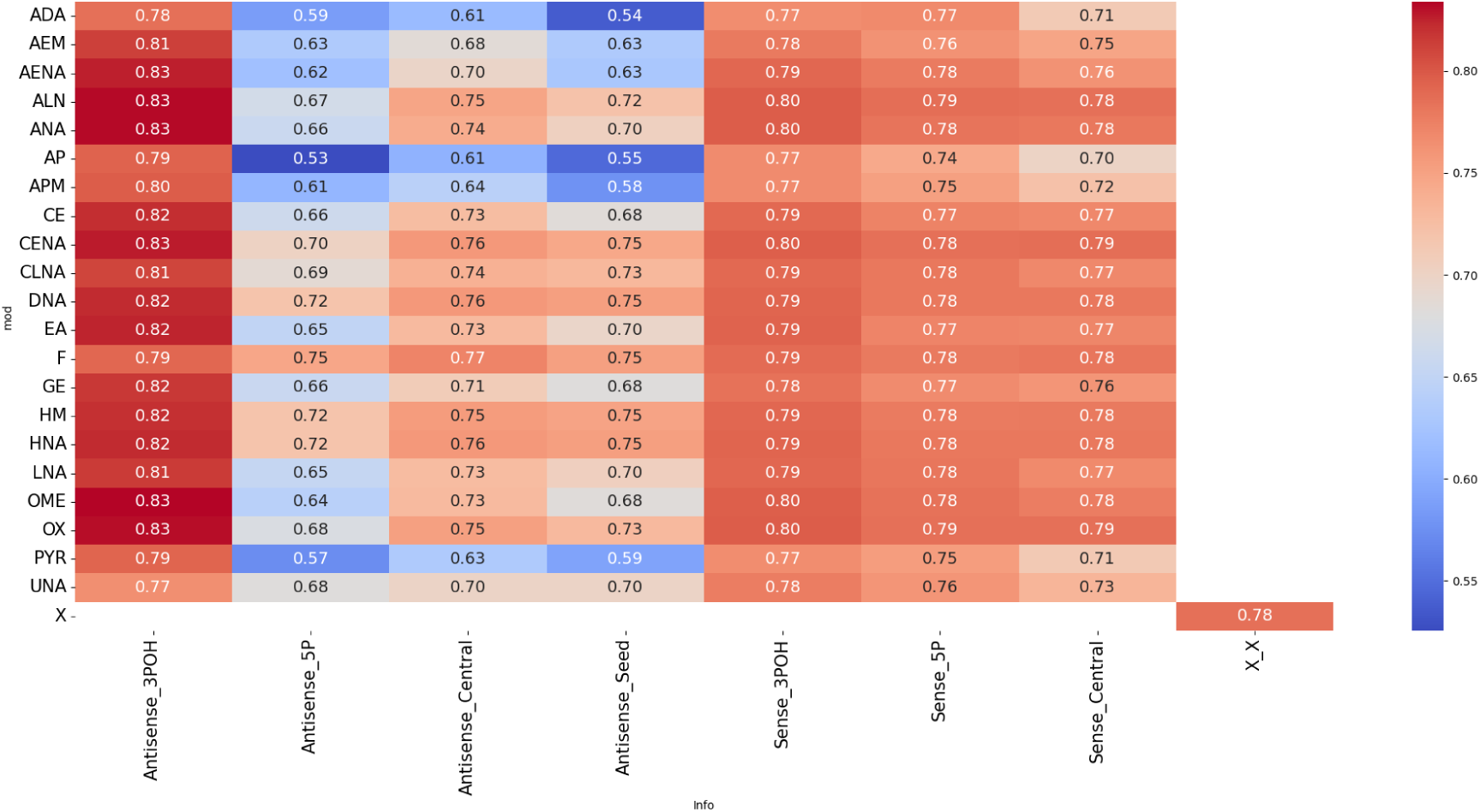
Design Patterns across generated dataset based on Bramsen Modifications are able to rediscover patterns within the dataset under different modification patterns.

### Ablation Study

To elucidate the contribution of each architectural component to overall model performance, we conducted a systematic ablation study on the cmsiRNA-mini which represents 10% of the cmsiRNAdb and Bramsen datasets, evaluating the significance of four key components: **Sequence**, **Chemistry**, **Cross-Attention**, **Metadata and Structure** and **MSE loss instead of Aleatoric**. To isolate the contribution of each component, we removed them one at a time from the full aleatoric model and re-evaluated performance. As shown in Figure 10, removing any single component from cmsiRNA-mini reduced performance, confirming that each modality contributes to the model’s predictive capability. This is expected, since the dataset spans diverse experimental setups, sequences, and chemistries. The pattern differs for Bramsen’s dataset in Figure 9, where the largest drop in AUC occurs when the chemistry component is removed. Because experimental conditions are uniform across this dataset, the metadata carries little or no information. The sequences are also largely consistent and provide minimal signal, so the cross-attention module contributes little as well. We therefore observe that the model performs as well as, or better than, the full model when these components are removed. We have also observed a drop in performance or widening of metric ranges across both the datasets when the loss function is changed to MSE instead of aleatoric, however noting that we also do not get the standard deviation prediction when using simple MSE loss.

**Figure 9:**
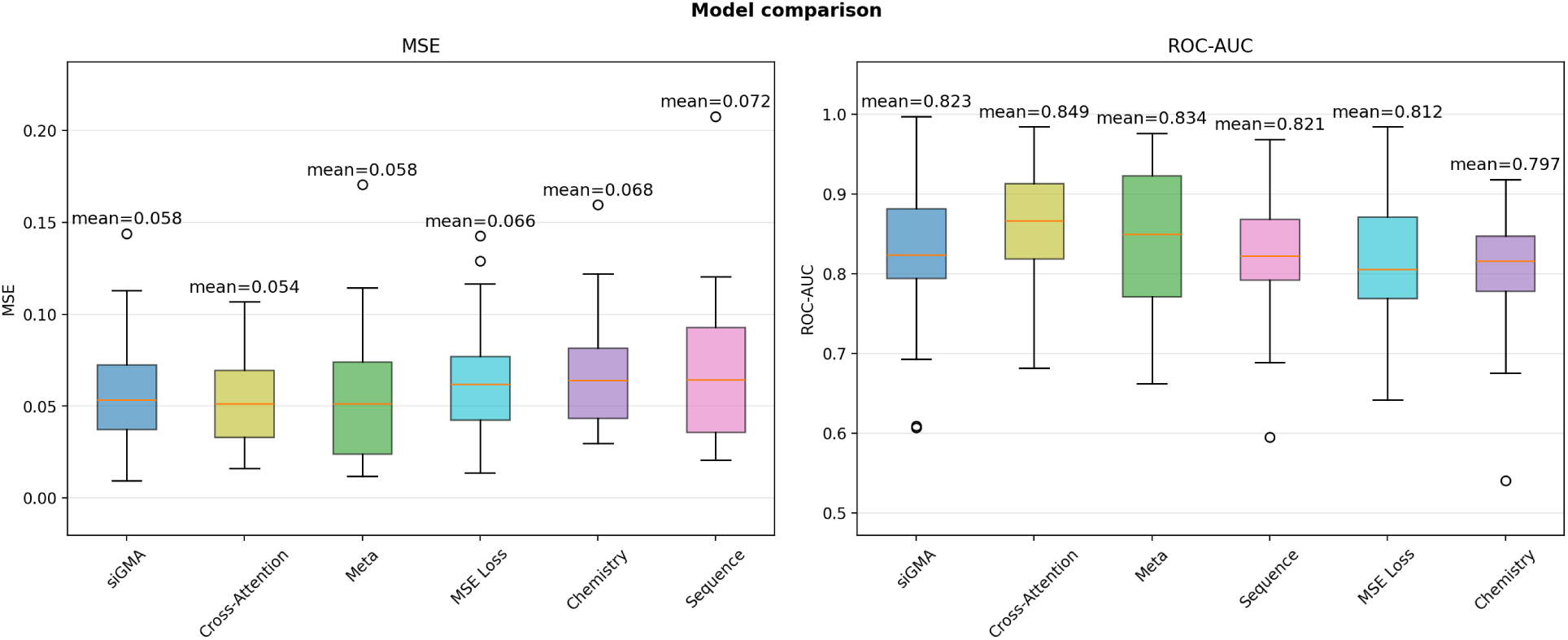
Ablation study on Bramsen Dataset shows a drop in performance with chemistry modality which is consistent with the data’s only variable signal.

**Figure 10:**
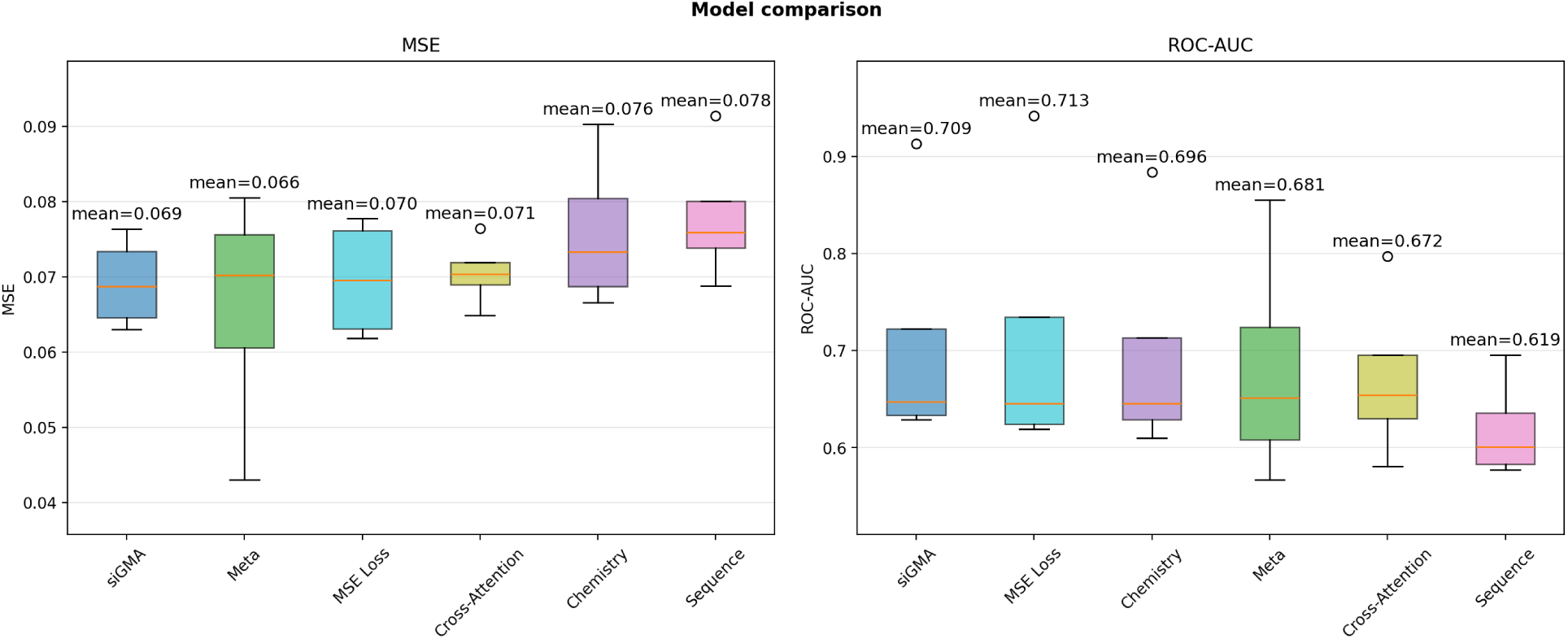
Ablation study on the cmsiRNA-mini shows a drop across all modalities due to variations available for each.

### Conclusion

In this study, we proposed siGMA, a heteroscedastic multimodal model, that leverages sequence, chemical modifications, experimental metadata and target accessibility scores and employs cross-attention to learn the complex sequence-chemistry-efficacy relationship underlying siRNA activity.

We highlight three key considerations, the first of which concerns the choice of cross-validation strategy. We observed substantial variability in model performance across groups, whether defined by biological target or by chemical modifications, and this heterogeneity is a critical dimension against which models should be evaluated. When this variability is not adequately accounted for, conventional approaches such as random k-fold cross-validation can produce overly optimistic and misleading performance estimates, as they fail to reflect how well a model generalizes to genuinely unseen targets or chemistry.

Secondly, the principal contribution of this work is the explicit modelling of noise. All datasets used here carry inherent label noise. The Huesken and Bramsen datasets are internally homogeneous with respect to experimental protocol, but the Bramsen dataset is sparsely populated across the chemical modification space, limiting the knowledge available to the model. The cmsiRNAdb, by contrast, provides approximately 40,000 data points, yet these are curated from several independent sources and therefore carry substantial label noise, arising chiefly from differences in knockdown measurement methods and assay choices. These conditions make it essential to model noise and uncertainty alongside point estimates, since point estimates alone may not faithfully reflect the underlying activity. Our results identify the origin of this uncertainty, which in most cases is attributable to inherent label noise rather than to model’s lack of examples. On this basis, we argue that noise modelling should be regarded as a necessary component of future AI-driven siRNA activity prediction.

Thirdly, we used the trained model to recover the design patterns encoded in the data, a necessary step before any model is applied to the design of new siRNAs. Establishing that the model rediscovers known sequence and chemistry preferences provides evidence that it has learned meaningful structure, and the patterns it surfaces may in turn suggest candidate designs that have not yet been tested experimentally.

Finally, although this work constitutes one of the earliest efforts to model noise in siRNA datasets, a number of avenues remain unexplored. We particularly stress the importance of systematic dataset generation for activity prediction, where chemistries, targets, and experimental conditions are deliberately and systematically varied under controlled settings. Considerable scope remains for more rigorous noise modelling in future work, especially in enriching experimental metadata and disentangling the individual contribution of each factor.

## AI Usage Declaration

The authors used generative AI tools during the preparation of this work. These tools were used to support tasks such as reviewing and refining written passages, improving clarity and grammar, and assisting with code development. All AI-generated content was reviewed, edited, and verified by the authors, who take full responsibility for the accuracy, integrity, and originality of the work.

## Author contributions statement

A.K. was responsible for Investigation and Formal analysis, Writing original draft and review & editing. A.M., A.W., G.K., A.L., and M.B. provided Supervision.

## References

(1) Alshaer, W.; Zureigat, H.; Al Karaki, A.; Al-Kadash, A.; Gharaibeh, L.; Hatmal, M. M.; Aljabali, A. A.; Awidi, A. European Journal of Pharmacology 2021, 905, 174178.

(2) Zhang, M.; Huang, Y. Trends in Molecular Medicine 2022, 28, 892–893.

(3) Saw, P. E.; Song, E.-W. Science China Life Sciences 2020, 63, 485–500.

(4) Khodagholi, Z.; Yousefi, N. NAR Genomics and Bioinformatics 2026, 8, lqag099.

(5) Martinelli, D. D. Health Sciences Review 2024, 11, 100157.

(6) Huesken, D.; Lange, J.; Mickanin, C.; Weiler, J.; Asselbergs, F.; Warner, J.; Meloon, B.; Engel, S.; Rosenberg, A.; Cohen, D.; Labow, M.; Reinhardt, M.; Natt, F.; Hall, J. Nature Biotechnology 2005, 23, 995–1001.

(7) La Rosa, M.; Fiannaca, A.; La Paglia, L.; Urso, A. International Journal of Molecular Sciences 2022, 23, 14211.

(8) Long, R.; Guo, Z.; Han, D.; Liu, B.; Yuan, X.; Chen, G.; Heng, P.-A.; Zhang, L. Briefings in Bioinformatics 2024, 25, bbae563.

(9) Bai, Y.; Zhong, H.; Wang, T.; Lu, Z. J. OligoFormer: an accurate and robust prediction method for siRNA design; preprint; Bioinformatics, 2024.

(10) Liu, B.; Yuan, Y.; Pan, X.; Shen, H.-B.; Jin, C. Med-X 2024, 2, 5.

(11) Liu, T.; Huang, J.; Luo, D.; Ren, L.; Ning, L.; Huang, J.; Lin, H.; Zhang, Y. International Journal of Biological Macromolecules 2024, 264, 130638.

(12) Tan, W.; Dai, M.; Ye, S.; Tang, X.; Jiang, D.; Chen, D.; Du, H. Journal of molecular biology 2025, 437, 169131.

(13) Chen, Y.; Tong, M.; Chen, L.; Li, W.; Zhu, K.; Li, J.; Tang, Y.; Liu, G. Journal of Medicinal Chemistry 2026, 69, 13434–13451.

(14) Karmakar, A.; Merii, A.; Weir, A.; Kudla, G.; Basham, M.; Lubbock, A. bioRxiv 2026, 2026–05.

(15) Dar, S. A.; Thakur, A.; Qureshi, A.; Kumar, M. Scientific reports 2016, 6, 20031.

(16) He, S.; Chen, C.; Pan, X.; Xue, G.; Yang, Y.; Feng, J.; Zulfiqar, H.; Zhang, Y.; Deng, K. BMC bioinformatics 2026, 27, 33.

(17) Hüllermeier, E.; Waegeman, W. Machine learning 2021, 110, 457–506.

(18) Salih, A. M.; Raisi-Estabragh, Z.; Galazzo, I. B.; Radeva, P.; Petersen, S. E.; Lekadir, K.; Menegaz, G. Advanced Intelligent Systems 2025, 7, 2400304.

(19) Bramsen, J. B.; Laursen, M. B.; Nielsen, A. F.; Hansen, T. B.; Bus, C.; Langkjær, N.; Babu, B. R.; Højland, T.; Abramov, M.; Van Aerschot, A., et al. Nucleic acids research 2009, 37, 2867–2881.

(20) Guo, F.-B.; Ou, H.-Y.; Zhang, C.-T. Nucleic acids research 2003, 31, 1780–1789.

(21) Bonidia, R. P.; Domingues, D. S.; Sanches, D. S.; De Carvalho, A. C. Briefings in bioinformatics 2022, 23, bbab434.

(22) Martinelli, D. D. Genomics 2024, 116, 110815.

(23) Adamczyk, J.; Ludynia, P. SoftwareX 2024, 28, 101944.

(24) Stiefl, N.; Watson, I. A.; Baumann, K.; Zaliani, A. Journal of chemical information and modeling 2006, 46, 208–220.

(25) Lorenz, R.; Bernhart, S. H.; Höner zu Siederdissen, C.; Tafer, H.; Flamm, C.; Stadler, P. F.; Hofacker, I. L. Algorithms for molecular biology 2011, 6, 26.

(26) Kendall, A.; Gal, Y. Advances in neural information processing systems 2017, 30.

(27) Lakshminarayanan, B.; Pritzel, A.; Blundell, C. Advances in neural information processing systems 2017, 30.

